# Characterizing Excretory-Secretory Products Proteome Across Larval Development Stages in *Ascaris suum*

**DOI:** 10.1101/2024.07.03.601870

**Authors:** Sergio Castañeda, Grace Adeniyi-Ipadeola, Yifan Wu, Charlie Suarez-Reyes, Antrix Jain, Juan David Ramírez, Jill E. Weatherhead

## Abstract

**Introduction:** *Ascaris lumbricoides* and *Ascaris suum* are parasitic nematodes that primarily infest the small intestines of humans and pigs, respectively. Ascariasis poses a significant threat to human health and swine health. Understanding *Ascaris* larval development is crucial for developing novel therapeutic interventions that will prevent ascariasis in both humans and pigs. This study aimed to characterize the excretory-secretory (ES) proteome of different *Ascaris suum* larval stages (L3-egg, L3-lung, L3-trachea) to identify potential targets for intervention to prevent *Ascaris*-induced global morbidity.

**Methods:** Stage-specific larvae were isolated, cultured in vitro and ES-product was collected. Third-stage *Ascaris* larvae (L3) were isolated from embryonated eggs (L3-egg), isolated from the lungs of Balb/c mice infected with *Ascaris suum* eggs at day 8 post infection (L3-lungs) and isolated from the trachea of Balb/c mice infected with *Ascaris suum* eggs at day 12 post infection (L3-trachea). ES products were obtained by culturing larvae. Proteomic analysis was conducted using liquid chromatography-tandem mass spectrometry (LC-MS/MS) and bioinformatic tools including MaxQuant, Perseus, and Andromeda, following a detailed protocol available on GitHub. The analysis encompassed peptide identification, scoring, and quantification against an organism-specific database, with subsequent quality control, correlation assessment, and differential abundance determination using the Amica algorithm.

**Results:** A total of 58 unique proteins were identified in the ES products. Fourteen proteins were common across all stages, while others were stage-specific. Principal component analysis revealed distinct protein profiles for each stage, suggesting qualitatively different proteomes. Gene ontology analysis indicated stage-specific GO enrichment of specific protein classes, such as nuclear proteins in L3-egg ES products and metabolic enzymes in L3-lung and L3-trachea ES products.

**Discussion:** This study revealed stage-specific differences in the composition of *Ascaris* ES products. Further investigation into the functional roles of these proteins and their interactions with host cells is crucial for developing novel therapeutic and diagnostic strategies against ascariasis.

## Introduction

*Ascaris lumbricoides* is a highly prevalent parasite globally [1]. According to the latest prevalence estimates there are approximately 800 million cases of ascariasis globally [2, 3]. Ascariasis disproportionately affects young children, with pre-school and school-aged children harboring the greatest worm burden [4–7]. In endemic regions, the use of anthelmintic preventive chemotherapy, through mass drug administration programs, aims to reduce worm burden and disease prevalence [2, 8, 9]. However, due to the ubiquitous contamination of *Ascaris* eggs in the environment, children endure recurrent infections throughout childhood [2, 9–12]. High worm burden and frequent re-infection of ascariasis results in a significant level of global morbidity, leading to nearly 754,000 Disability Adjusted Life Years (DALYs) [9, 13]. As a result of *Ascaris-*induced long-term morbidity, there is an urgent need to develop novel therapeutic interventions, like vaccines, to prevent ascariasis. However, developing therapeutic interventions that prevent a high burden of disease and reinfection will require an in-depth understanding of how *Ascaris* larvae develop into adult worms and how they interact with the host immune response [14]. Similarly, *Ascaris suum* presents significant health and economic challenges to the swine industry due to its widespread infection among pigs worldwide [1, 15]. *A. suum* can affect pigs of all age groups, but its prevalence within specific age categories, particularly young pigs, is often influenced by housing conditions and management practices. Overall ascariasis in pig populations leads to significant economic losses due to reduced fee conversion efficiency and increased condemnation resulting in production losses [16, 17].

*Ascaris lumbricoides* and *Ascaris suum* are nearly identical morphologically and genetically and routinely cause cross-infection between hosts [1, 18, 19]. Both *Ascaris* spp. larvae undergo an essential migratory phase that promotes development from larvae to mature adult worm [20]. What distinguishes the life cycle of *Ascaris* spp. from other soil-transmitted helminths is its hepato-tracheal migration. Following ingestion of embryonated *Ascaris* eggs from the environment, third-stage larvae (L3) emerge from the eggs in the intestines. L3 migrate from the intestines to the liver, then proceeds to the lungs via the circulatory system. Within the lung parenchyma, L3 larvae develop into mature L3 as they are ascending the trachea and swallowed back into the intestines. In the intestines, L3 mature first into L4 larvae and than into adult worms [20, 21].

Throughout its migration, *Ascaris* spp larvae release a wide range of molecules termed excretory-secretory (ES) products. These products encompass antimicrobial peptides, proteins, lipids, metabolites, among others, and are integral in shaping the intricate interactions between the host and the parasite. They influence key aspects such as worm survival, migration, and immune modulation, ultimately aiding in the successful establishment and persistence of the parasite within the host [22–25].

With significant advancements in mass spectrometry and genomic technologies, numerous challenges and constraints in the proteomic analysis of parasite ES proteins have been effectively addressed. Consequently, ES proteomes for parasitic nematodes such as *Ancylostoma caninum*, *Brugia malayi*, *Haemonchus contortus*, *Teladorsagia circumcincta*, and *Trichinella spiralis* have been successfully characterized [26–32]. Unraveling the composition and functional significance of ES products across *Ascaris* life cycle presents a compelling avenue to advance our understanding of parasite biology and potential targets for intervention strategies.

Given the close relation between *A. suum* and *A. lumbricoides*, and the availability of the *A. suum* genome, we undertook the characterization of the protein composition of ES in three distinct larval stages of *Ascaris* (L3-egg, L3-lung, and L3-trachea) using tandem mass spectrometry annotated with the *A. suum* genome in a mouse model. Identifying highly abundant and conserved, stage-specific proteins within the *Ascaris* ES proteome presents a promising avenue for discovering novel targets that could be utilized in the development of preventive and therapeutic interventions against *Ascaris* infection.

## Methods

### Stage-specific larval isolation

#### Hatching of third-stage larvae (L3 egg stage)

Embryonated eggs in H_2_SO_4_ solution were centrifuged, the supernatant was removed, and the pellet was re-suspended in sodium hypochlorite solution and incubated for 2 hours. After incubation, the eggs were centrifuged with subsequent removal of the supernatant. The eggs were washed with ultra-pure water 3 times and re-suspended in Hanks Balanced Salt Solution (HBSS) with pH 2.0 using hydrogen chloride to achieve the desired pH and incubated for 30 minutes in 37°C + 5% CO_2_. The eggs were centrifuged, the supernatant was removed while the pellet was re-suspended in 10 mL of HBSS solution pH 7.0 to neutralize eggs for 30 minutes. The eggs were centrifuged, and the egg pellet was re-suspended in culture media (RPMI-1640) with penicillin-streptomycin (4% pen/strep). The egg solution was placed in 6-well culture plate (4mL per well) and incubated over 96 hours. Egg hatching was monitored using light microscopy. Once the larvae hatched, new media was placed in the culture plates and incubated overnight at 37°C +5% for 48 hours. The experiment was completed in triplicate.

#### Collection of third stage larvae from the lungs (L3-lung)

5 Balb/c mice were infected with 2500 *Ascaris suum* eggs in a total volume of 100 μL distilled water by oral gavage. After 8 days of infection, mice were euthanized by intraperitoneal injection of ketamine (up to 150 mg/kg) and xylazine (up to 15 mg/kg). After euthanasia, the lungs were removed, macerated in a petri dish, placed in a modified Baermann apparatus in warm PBS and incubated at 37°C +5% CO_2_ for 4 hours. The fluid in the Baermann apparatus was transferred to a 50 mL conical tube and centrifuged. The supernatant was removed, and the pellet was washed with ultra-pure water three times. After the final wash and removal of supernatant, the larvae were transferred into a 6-well culture plate with 4 mL of culture media to each well. The culture plates were incubated at 37°C +5% CO_2_ for 48 hours. The experiment was completed in triplicate.

#### Collection of fourth-stage larvae from the trachea (L3 trachea)

5 Balb/c mice were infected as described above. On post-infection day 12, the mice were euthanized, and a tracheostomy was created by placing a 20-gauge angiocatheter into the trachea. The angiocatheter was flushed with 0.8 mL of PBS three times, all returned fluid was collected and passed through a 40µm cell strainer. The 40µm cell strainers were washed using 4mL of PBS to release the larvae into a 6 well cell culture plate. Culture plates were rested for 5 minutes to allow larvae to settle to bottom of the culture plate with gravity. PBS was removed from the wells. The larvae were then washed with 4mL distilled water a total of three times. After completion of the wash, 4 mL of culture media was added to each well and the 6-well culture plates were incubated at 37°C +5% CO_2_ for 48 hours. The experiment was completed in triplicate.

### Preparation and proteomic analysis of ES products

After 48 hours the supernatant was pooled, filtered through a 0.45 µm filter (PALL Corporation), frozen in liquid nitrogen and stored until use as ES product. This process was repeated in three separate experiments. Using mass spectrometry at the Baylor College of Medicine Mass Spectrometry Proteomics core a total of 40ml volume of ES product from each developmental stage was lyophilized and trypsin digested using S-Trap™ (Protifi, NY) according to manufacturer’s protocol. Digested peptides were cleaned using a C18 disk plug (3M Empore C18) and dried in a speed vac. The peptide concentration was normalized to a total of 1 ug prior to the mass spectrometry run. Liquid chromatography-tandem mass spectrometric (LC-MS/MS) analysis was carried out using a nano-LC 1200 system (Thermo Fisher Scientific, San Jose, CA) coupled to Orbitrap Fusion™ Lumos ETD mass spectrometer (Thermo Fisher Scientific, San Jose, CA). Peptides were loaded on a C-18 trap-column (Reprosil-Pur Basic C18, 2cm X 100µm, ID 1.9 µm, Dr. Maisch GmbH, Germany) and eluted using a 75min gradient of 5-28% acetonitrile/0.1% formic acid at a flow rate of 750nl/min on a C-18 analytical column (Reprosil-Pur Basic C18, 5cm x 150µm, ID 1.9 µm, Dr. Maisch GmbH, Germany). The peptides were directly electro-sprayed into the mass spectrometer operated in a data-dependent mode with ‘top 30’ method. The full MS scan was acquired in Orbitrap in the range of 300-1400m/z at 120,000 resolution followed by MS2 in Ion Trap (HCD 28% collision energy) with 15sec dynamic exclusion time.

### Bioinformatic analysis of ES products

The analytical framework employed MaxQuant, Perseus, and Andromeda, as detailed at https://github.com/FredHutch/maxquant-pipeline. This protocol encompassed the identification of peptides, determination of mass, and intensity of peptide peaks in mass spectrometry (MS) spectra, thereby facilitating the identification of isotopic patterns [33–35].

Search parameters were a precursor tolerance of 20 ppm, MS/MS tolerance of 0.05 Da, Dynamic modification of Oxidation and protein N-terminal Acetylation was allowed. The query of peptide and fragment masses was conducted against a database housing organism-specific sequences of *Ascaris*, obtained from UniProt (https://www.uniprot.org/uniprotkb?facets=reviewed:false&query=(taxonomy_id:6253)) PMID:21685128. Subsequently, scoring through a probability-based method termed peptide scoring was performed, with the implementation of a false discovery rate (FDR) approach to limit the occurrence of matches by chance. FDR was ascertained through statistically rigorous methodologies addressing multiple hypothesis testing. Additionally, the database search incorporated not only target sequences but also their reverse and contaminant counterparts, aiding in the establishment of a statistical cutoff for acceptable spectral matches. Peptides were accepted based on a false discovery rate (FDR) of <0.05 at both the peptide and protein levels. Proteins were quantified using the LFQ value from MaxQuant employing default settings [33–35].

Following this preprocessing, the acquired outputs are undergoing a comprehensive analysis encompassing quality control (QC), correlation assessment, density analysis, differential abundance determination, and group comparisons, facilitated by the Amica algorithm (https://github.com/tbaccata/amica) [36], using default parameters: Log2 fold change threshold 1.5; significance cutoff (which value to use) 0.05.

To perform gene ontology analysis, the software QuickGO and PROTEINSIDE was used to determine GO Biological Process and Cellular Component ontology enrichments using the *Ascaris suum* genome as a reference (PRJNA62057). Additionally, DeepLoc was employed to identify proteins with signal peptides within the dataset, thereby identifying secreted proteins [37].

Subsequent analyses of the output obtained by MaxQuant were evaluated also by Perseus [33–35]. Finally, we implement the open-source tool Protter, for visualization of proteoforms and interactive integration of annotated and predicted sequence features [38].

## Results

### Protein identification

The proteins within the ES products of different *Ascaris* larval stages (L3-egg, L3-lung, L3-trachea) were characterized using LC-MS/MS. MaxQuant searches of the data identified 58 unique proteins by Gene ID in the ES products of all three larval developmental stages combined (Table 1).

**Table 1.**
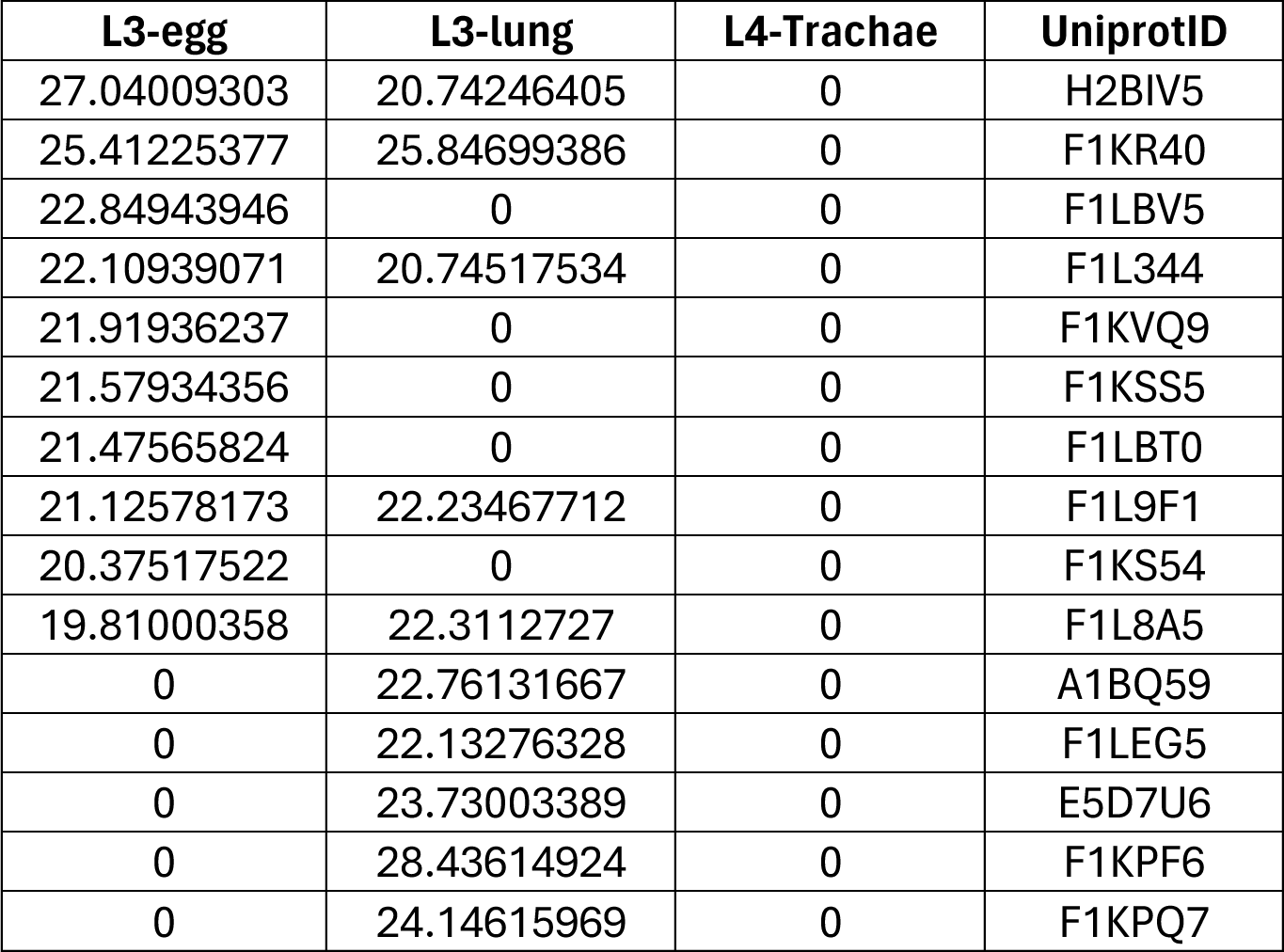
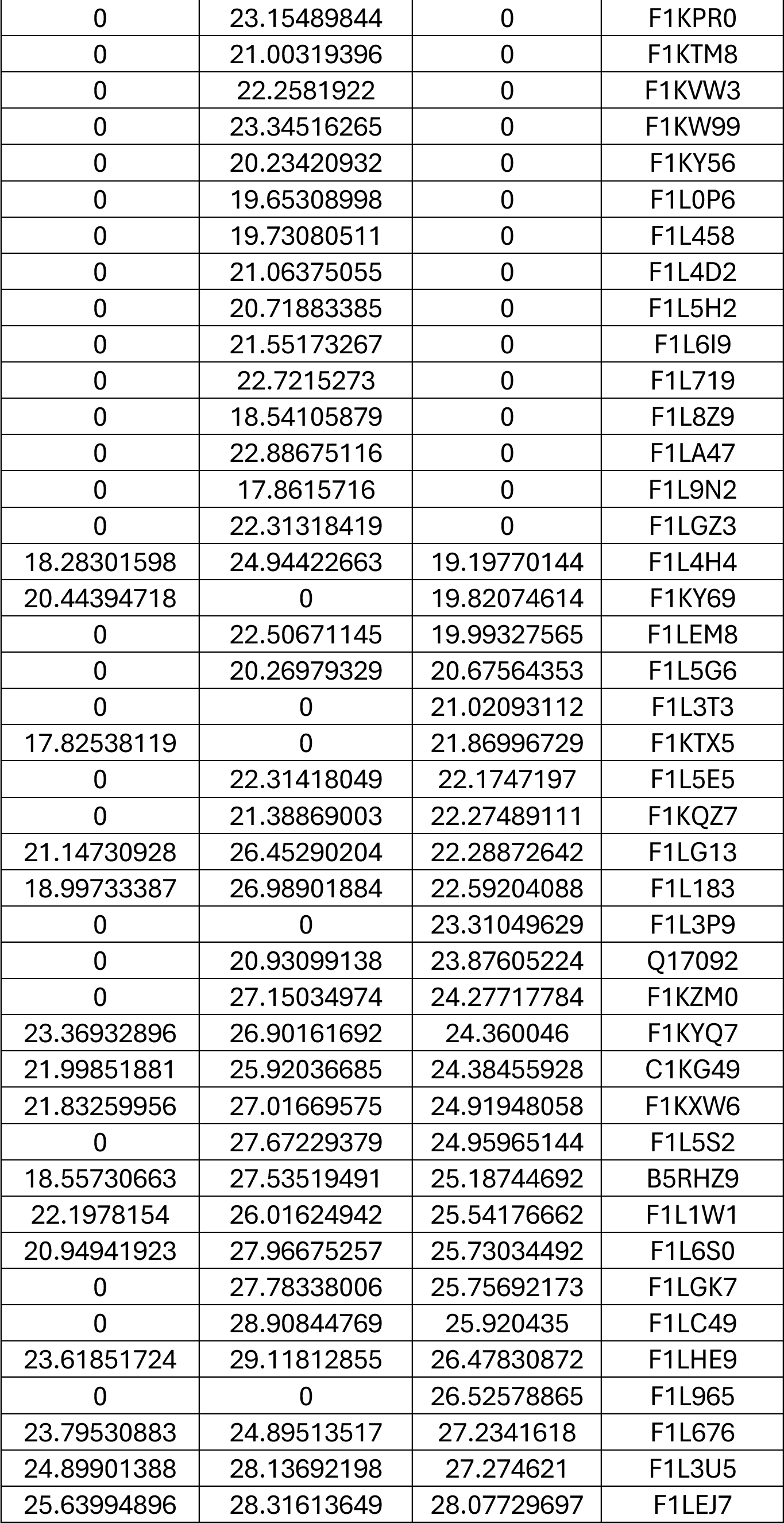
Quantification values for each protein were calculated using the LFQ values obtained from MaxQuant at each larval stage.

Specifically, 5 proteins were exclusively identified in L3-egg, while L3-lung exhibited 20 unique proteins, and L3-trachea displayed 3 proteins exclusive to its stage. Additionally, 14 proteins were detected in ES products conserved across all three larval development stages (Figure 1a). The protein profiles of the ES products from each larval developmental stage were compared using principal component analysis (PCA) to assess their relatedness. The PCA revealed three distinct proteome clusters corresponding to each stage (Figure 1b).

**Figure 1.**
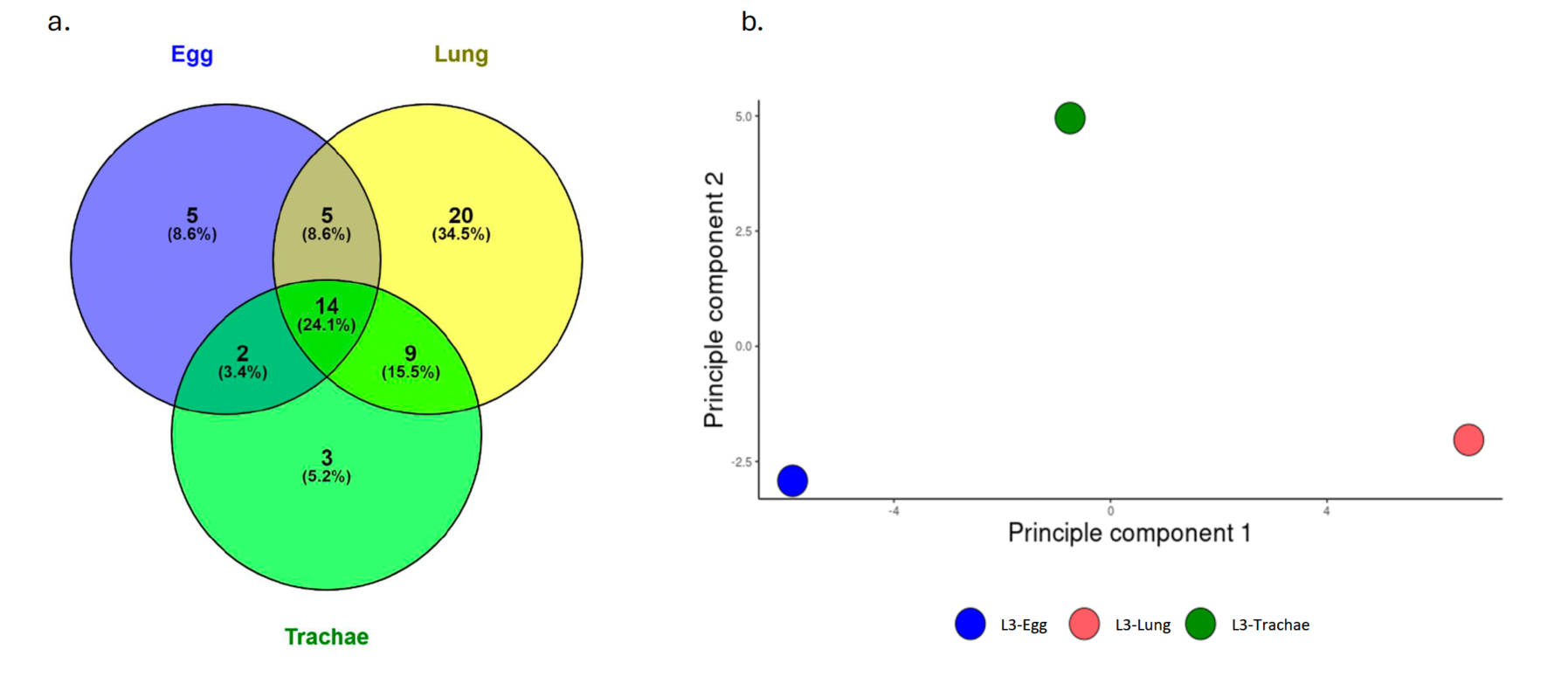
(a) The Venn diagram illustrates the distribution of unique and shared proteins across the ES product of *Ascaris* stages. (b) Principal Component Analysis (PCA) reveals that the overall composition remains distinctive for each developmental stage ES product.

### Comparative analysis

We further compared the proteomic profiles between different larval stages using volcano plots and heat maps. It is noteworthy that, although statistically significant differences were not observed among the analyzed samples, distinct groups of proteins associated with L3-egg, L3-lung, and L3-Ttachea ES products were identified, alongside some shared proteins (Figure 2a). The lack of difference in protein enrichment between stages’ ES products may be attributed to the similarity between the proteomes or to an insufficient number of proteins for a robust comparison due to the heterogeneity in the samples. Additionally, based on the obtained results, Spearman’s correlation analysis reveals distinct correlation coefficients among the various protein groups across stages. Specifically, a correlation of 0.54 is observed between L3-lung and L3-trachea ES proteins. In contrast, the correlation between L3-egg and L3-lung ES proteins is determined to be 0.21, while the correlation between L3-egg and L3-trachea ES proteins is calculated to be 0.32 (Figure 2b).

**Figure 2.**
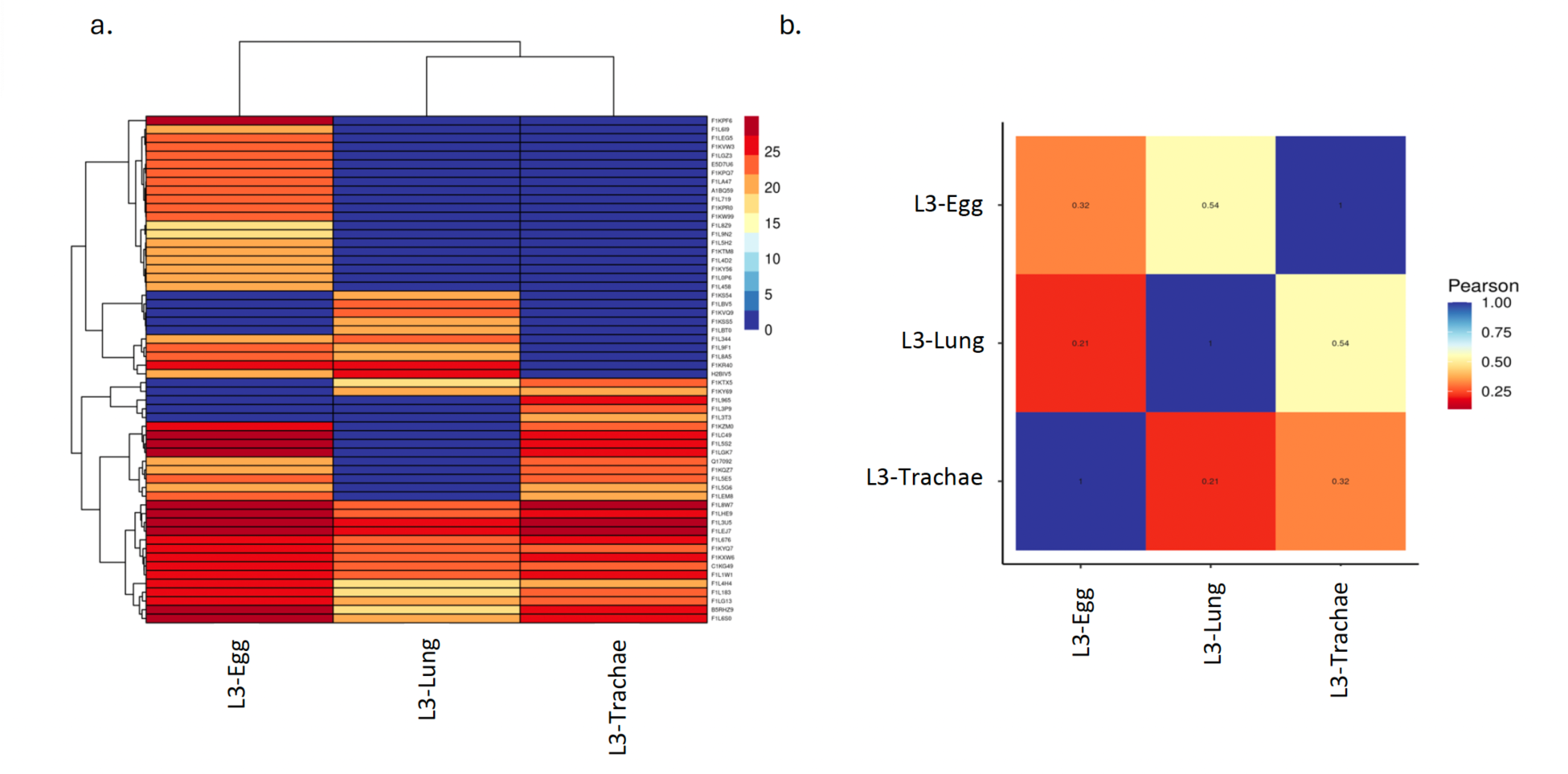
(a) HeatMap of protein profile identified by each developmental ES product stage of *Ascaris*. (b) Spearman’s correlation analysis among the *Ascaris*-related protein groups.

### Gene ontology

The Gene Ontology (GO) annotation of proteins enriched in the ES product was analyzed by molecular function, cellular component, and biological process (Table 2, Figure S1), as well as according to their property of being secreted proteins (Table S1).

**Table 2.**
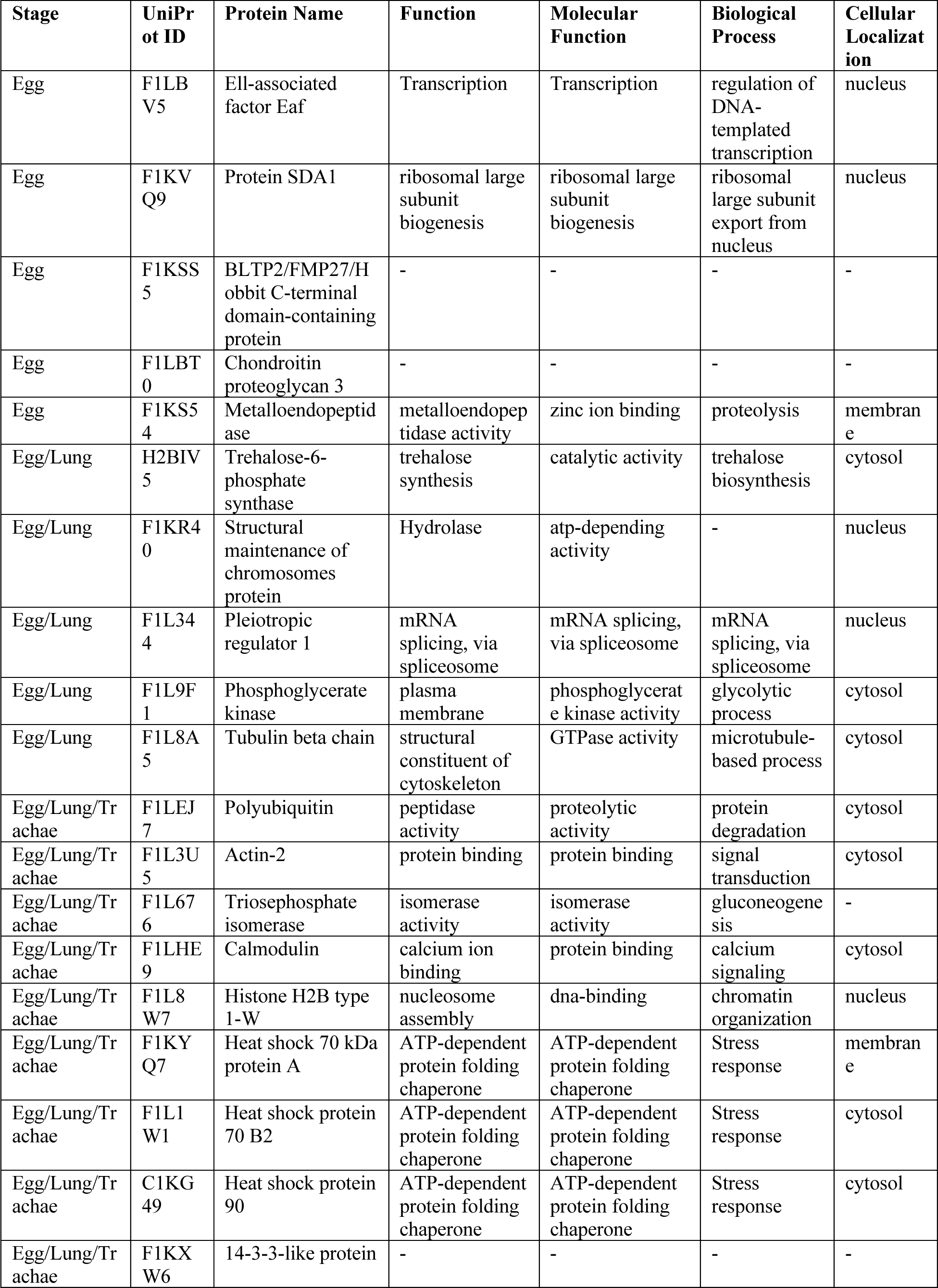
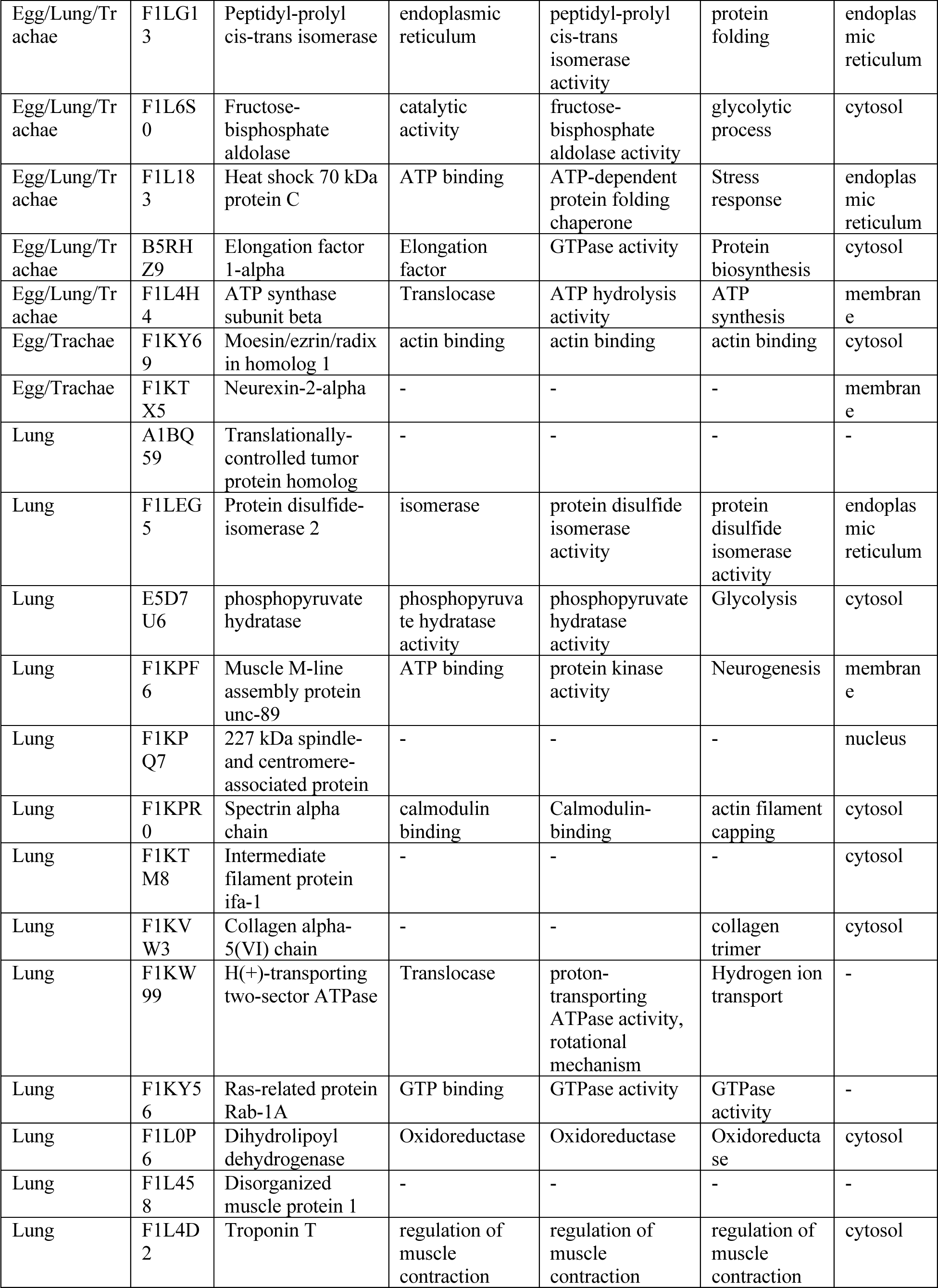
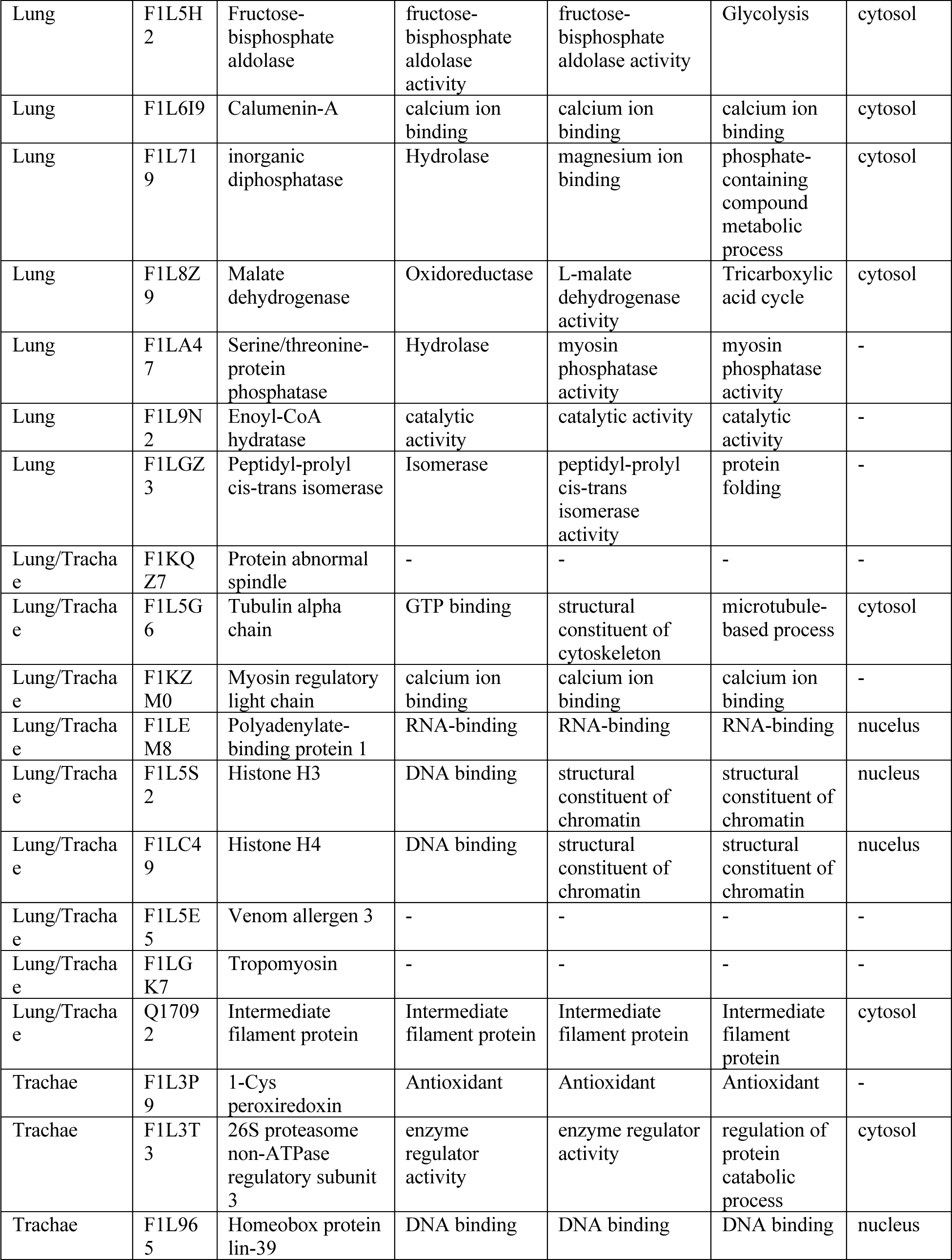
Identified protein function, implicated biological processes, and subcellular localization.

Additionally, based on annotated and predicted protein sequences derived from previous analyses, we conducted a visual reconstruction of proteoforms of the main proteins identified. This aimed to represent protein sequence characteristics within the context of protein topology and experimental proteomic evidence (Figure 3). Notably, proteins identified in L3-egg ES product exhibit a pronounced association with nuclear proteins engaged in replicative and transcriptional processes, including Ell-associated factor Eaf, Protein SDA1, BLTP2/FMP27/Hobbit C-terminal domain-containing protein, Chondroitin proteoglycan 3, and Metalloendopeptidase. Conversely, proteins identified in L3-lung and L3-trachea ES products predominantly pertain to cytoplasmic proteins involved in energetic processes, such as glycolysis, and proteins related to the cytoskeleton (Figure S2, S3). For L3-lung ES products, the identified proteins include translationally-controlled tumor protein homolog, Protein disulfide-isomerase 2, phosphopyruvate hydratase, Muscle M-line assembly protein unc-89, 227 kDa spindle-and centromere-associated protein, Spectrin alpha chain, Intermediate filament protein ifa-1, Collagen alpha-5(VI) chain, H(+)-transporting two-sector ATPase, Ras-related protein Rab-1A, Dihydrolipoyl dehydrogenase, Disorganized muscle protein 1, Troponin T, Fructose-bisphosphate aldolase, Calumenin-A, inorganic diphosphatase, Malate dehydrogenase, Serine/threonine-protein phosphatase, Enoyl-CoA hydratase, and Peptidyl-prolyl cis-trans isomerase (Figure S2, S3). On the other hand, for L3-trachea ES products, proteins found include 1-Cys peroxiredoxin, 26S proteasome non-ATPase regulatory subunit 3, and Homeobox protein lin-39. These findings underscore distinct molecular emphases and functional roles across different developmental stages of *Ascaris*, shedding light on the nuanced biological processes characterizing each stage.

**Figure 3.**
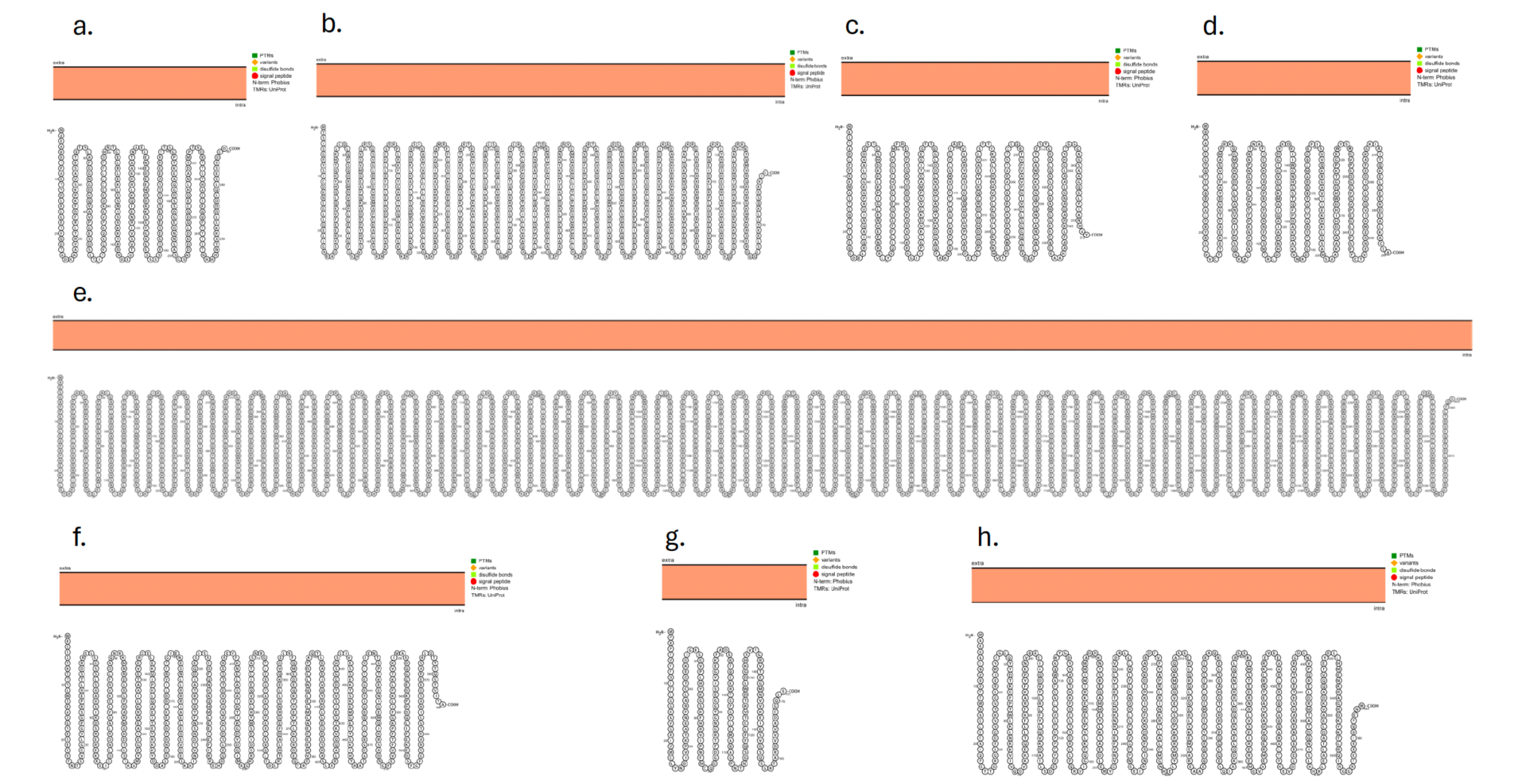
Visualization of proteoforms and interactive integration of annotated and predicted sequence features of main proteins. Nuclear proteins associated with replication and transcription, such as (a) Ell-associated factor Eaf (UniProt ID F1LBV5) and (b) Protein SDA1 (UniProt ID F1KVQ9) in L3-egg ES product. Glycolytic enzymes like (c) fructose-bisphosphate aldolase (UniProt ID F1L5H2) and (d) malate dehydrogenase (UniProt ID F1L8Z9) in L3-lung ESP. Cytoskeletal proteins such as (e) spectrin alpha chain (UniProt ID F1KPR0) and (f) intermediate filament protein ifa-1 (UniProt ID F1KTM8) in both L3-lung and L3-trachea ES product. (g) Peptidyl-prolyl cis-trans isomerase (UniProt ID F1LGZ3), an immunophilin found in L3-Lung ES product. Identified (h) Ezrin-radixin-moesin (ERM) proteins (UniProt ID F1KY69) in the secreted proteomes of larval development stages L3-lung and L3-trachea ES product.

## Discussion

This study employed LC-MS/MS to characterize the protein profiles of excretory-secretory (ES) products from different *Ascaris* larval stages (L3-egg, L3-lung, L3-trachea). This is the most comprehensive evaluation of stage-specific ES proteomes during *Ascaris* larval migration to date. Findings reveal intriguing insights into the stage-specific composition of these secretions, potentially reflecting distinct biological functions during development. The migration of *Ascaris* larvae through the host is a crucial developmental process for the parasite. This study elucidates the similarities and differences in the secreted proteome among L3 developmental stages, offering valuable insights that could spur research focused on identifying novel diagnostic and vaccine targets essential for controlling and eradicating ascariasis [22, 39–41].

Prior research has investigated ES proteins derived from adult worms, uncovering a diverse array of proteins, enzymes, and biomolecules with roles in immunomodulation, immunoevasion, and tissue degradation. Among these are Glycosyl hydrolases, crucial for breaking down complex carbohydrates and integral to the parasite’s energy metabolism. Additionally, C-type lectins exhibit a wide range of functions including cell-cell adhesion, immune responses to pathogens, and involvement in apoptosis. Furthermore, Heat shock proteins (HSPs) have been identified, which may modulate host responses to facilitate and sustain infection, among other functions. [22, 40–43]. Furthermore, a study evaluating ES protein of *Ascaris* L3-egg, L3-lung and L4-intestines was completed in two pigs. The majority of the proteins identified were stage-specific but 14-3-3-like protein and a serpin-like protein were conserved amongst the three different larval stages. While pigs are a natural host of ascariasis, use of a pig model is limiting due to size and cost [26]. Here, we used a highly replicated mouse model of ascariasis in order to repeat the experiment in triplicate and identify variation between each larval development stage as well as within each larval development stage.

The identification of stage-specific proteins in our study aligns with previous observations in other parasitic nematodes, where ES products exhibit dynamic changes throughout the life cycle [27, 43–48]. For instance, in *Trichinella spiralis*, infective larvae secrete proteins like excretory-secretory antigen-like protein (TES) and cystatin inhibitor during muscle invasion, potentially aiding tissue penetration and immune modulation [49, 50]. Similarly, *Angiostrongylus cantonensis* ES products change significantly between different developmental stages, with glutathione S-transferase and superoxide dismutase enriched in adult stages, suggesting roles in detoxification and stress management [51, 52].

In the current analysis, a total of 58 unique proteins were identified in the *Ascaris* ES products. Notably, 5 proteins were exclusively identified in L3-egg ES product, 20 proteins unique to proteins were unique to L3-lung ES product, and 3 proteins were distinct to L3-trachea ES product. Additionally, 14 detected proteins were conserved across all three larval developmental stages (Figure 1a, 2a). Protein diversity and protein abundance may be secondary to the active migration. In comparison to the intraluminal L3 egg stage, the L3-trachea stage and L3 lung stage, which require migration through tissue and molting, may necessitate upregulation and secretion of proteins to facilitate these metabolically taxing processes [21, 27, 53, 54].

Our research into L3-egg ES product supports the idea that the GO enrichment of nuclear proteins associated with replication and transcription, such as Ell-associated factor Eaf (UniProt ID F1LBV5) and Protein SDA1 (UniProt ID F1KVQ9), correlates with active cell division and differentiation during this early developmental stage (Figure 2, 3). This finding aligns with previous studies on *Ascaris suum* embryogenesis, where similar proteins were identified [27, 41, 53]. Thus, it appears that ES products in L3-egg may play a crucial role in establishing the foundation for further larval development. Conversely, in L3-lung and L3-trachea ES products, we observed a dominance of cytoplasmic proteins related to energy metabolism and cytoskeletal functions, which aligns well with the increased demands of these later stages [27, 41, 53]. For example, the presence of glycolytic enzymes like fructose-bisphosphate aldolase (UniProt ID F1L5H2) and malate dehydrogenase (UniProt ID F1L8Z9) in L3-lung ES product suggests an adaptation to anaerobic environments during lung migration (Figure 2, 3) [55, 56]. Moreover, the detection of cytoskeletal proteins such as spectrin alpha chain (UniProt ID F1KPR0) and intermediate filament protein ifa-1 (UniProt ID F1KTM8) in both L3-lung and L3-trachea ES product suggests their role in preserving cell shape and aiding in active movement within the host, as has been observed in various nematode models, including *Trichinella britovi* and *Caenorhabditis elegans* (Figure 2, 3) [57–59]. These findings underscore the potential significance of ES products in facilitating the heightened metabolic activity and motility crucial for the successful migration and establishment of larvae within the host. Further exploration of these specific proteins could yield therapeutic and diagnostic targets for *Ascaris* larval migration, offering valuable insights into proteins essential for helminthic metabolic and motility processes vital to *Ascaris* development.

Furthermore, our analysis revealed several interesting proteins within the ES products specific to particular larval stages. For instance, peptidyl-prolyl cis-trans isomerase (UniProt ID F1LGZ3), an immunophilin found in L3-lung ES product, exhibits activities that isomerize peptidyl-proline bonds, thereby facilitating enhanced protein folding (Figure 2, 3) [60, 61]. Immunophilins, major binding proteins for certain immunosuppressive drugs, have been found in other helminth ES products, including *Trichinella spiralis* early larval development stages [62]. While detailed functional analysis is available for many protozoal immunophilins, relatively little is known about the functionality of helminth immunophilins outside of *Caenorhabditis elegans*, although they are widely expressed within helminth species [61]. Parasite-derived immunophilins could serve as potential targets for diagnostic assays or drug development, especially when considering non-immunosuppressive analogs. Cyclosporin A, along with its non-immunosuppressive counterparts, has demonstrated inhibitory effects on parasites, suggesting promising avenues for therapeutic intervention [60, 61].

Additionally, we identified Ezrin-radixin-moesin (ERM) proteins (UniProt ID F1KY69) in the secreted proteomes of larval development stages L3-lung and L3-trachea ES proteins (Figure 2, 3). ERMs are proteins at the intersection of the cytoskeleton and the extracellular membrane, traditionally considered structural proteins allowing cross-linking of actin to the apical cellular membrane [63]. However, recent studies have revealed significant roles for ERMs in metabolic processes, cell signaling, and activation of the immune response [63, 64]. While several studies have emphasized the role of ERM in protozoa infectivity, particularly in *Trypanosoma cruzi*, there is currently no indication of the involvement of ERM in helminth larval migration in the context of ascariasis [65]. It has been identified that ERM proteins could be associated with various parasite processes, including invasion, potentially regulating F-actin dynamics and plasma membrane interplay. Additionally, their involvement in the development of several structures has been observed in models such as *C. elegans* [66]. Furthermore, changes in the phosphorylation state of these proteins may be linked to interactions with specific tissues, offering valuable insights into larval changes associated with specific anatomical and parasite-host-tissue interactions during larval migration [63–67].

Although statistically significant differences in protein abundance between stages were not detected, several lines of evidence suggest qualitative differences in the ES proteomes. First, principal component analysis (PCA) revealed distinct clusters for each stage, indicating unique protein profiles (Figure 1b). For instance, L3-egg ES product clustered separately due to the presence of embryonic development proteins like Eaf and SDA1, absent in later stages [39, 41]. Conversely, L3-lung and L3-trachea ES product formed a separate cluster enriched in metabolic enzymes like aldolase and malate dehydrogenase, suggesting a shared focus on energy production. Second, the identification of unique proteins in each stage further supports qualitative differences. For example, L3-egg ES product exclusively contained chromatin remodeling proteins, essential for early development, while L3-lung ES product exhibited specific cytoskeletal proteins like spectrin and intermediate filament proteins, potentially involved in motility during lung migration [55, 56]. Additionally, L3-trachea ES product uniquely possessed stress response proteins like peroxiredoxin, suggesting adaptation to the host environment [55, 56]. Third, the heterogeneity of ES products and inherent variability could mask subtle abundance differences between stages, making statistical detection challenging. For instance, the presence of immunomodulatory proteins in all stages might mask stage-specific variations in their abundance, requiring targeted analyses for accurate comparisons. It is important to highlight the relatively small number of identified proteins might not capture the full complexity of the proteomes [68–72]. Focusing on a broader range of proteins or employing targeted approaches might reveal significant differences between stages. Therefore, while statistically significant differences in protein abundance weren’t observed, the distinct clustering, stage-specific protein groups, and potential limitations suggest qualitative differences in the proteomes of *Ascaris* ES products across developmental stages. Further studies with larger sample sizes and targeted protein analyses are crucial to fully understand these qualitative variations and their functional implications.

These findings pave the way for further investigations into the specific functions of individual proteins and their contributions to the overall biological processes of each larval stage. Studying the downstream effects of Ell-associated factor Eaf through gene expression analysis could elucidate its role in regulating cell cycle genes in L3-egg [39, 41]. While nzyme activity assays could measure the specific activity of fructose-bisphosphate aldolase and malate dehydrogenase in different stages, providing direct evidence for their contribution to energy production [56]. Furthermore, for instance, immunofluorescence microscopy could visualize the localization of spectrin alpha chain and intermediate filament protein ifa-1 in migrating larvae, revealing their role in maintaining specific cellular structures and facilitating movement [58, 59]. Beyond individual proteins, exploring the interactions between ES products and host cells could provide valuable insights into the mechanisms underlying *Ascaris* pathogenesis. For example, investigating how cytokines secreted by host immune cells in response to specific ES proteins could reveal their immunomodulatory potential. Additionally, studying the binding of ES proteins to host cell receptors could identify potential targets for therapeutic intervention [27, 51, 52, 73, 74]. While further investigations are necessary to fully elucidate the functional significance of these proteins and their interactions with host cells, these findings contribute to a deeper understanding of the complex biology of *Ascaris* and its adaptations to the parasitic life cycle.

## Author contributions

SC, GAI, YFW, JDR, JW conceived and designed experiments. GAI, CSR, YFW performed laboratory experiments or clinical sampling. SC, AJ, JDR provided technical support and assisted with data collection. SC, AJ, performed data analysis. SC, GAI, YW, JDR, JW wrote the manuscript. All authors contributed to reviewing the paper and agreed on the present version for submission.

## Data availability

Preprocessing and processing results data are available at https://github.com/scastanedag/Proteome_Larval_Development-Stages_Ascaris_suumv

## Supplementary material

**Figure S1.** ES product proteins and their corresponding subcellular locations at each larval stage.

**Figure S2.** GO enrichment analysis by Cellular Component. It shows significant results obtained in a. L3-egg and L3-lung.

**Figure S3.** GO enrichment analysis by Biological Process. It shows significant results obtained in a. L3-egg and L3-lung.

**Table S1.** Secreted proteins identified. The Predicted localizations and Predicted signals display the subcellular localizations and sorting signals predicted for the query protein, respectively. The Probability table displays the probability assigned by the model to each of the subcellular localizations.

