## Supplementary figures and images for "Characterizing Excretory-Secretory Products Proteome Across Larval Development Stages in *Ascaris suum*"

### Figure S1

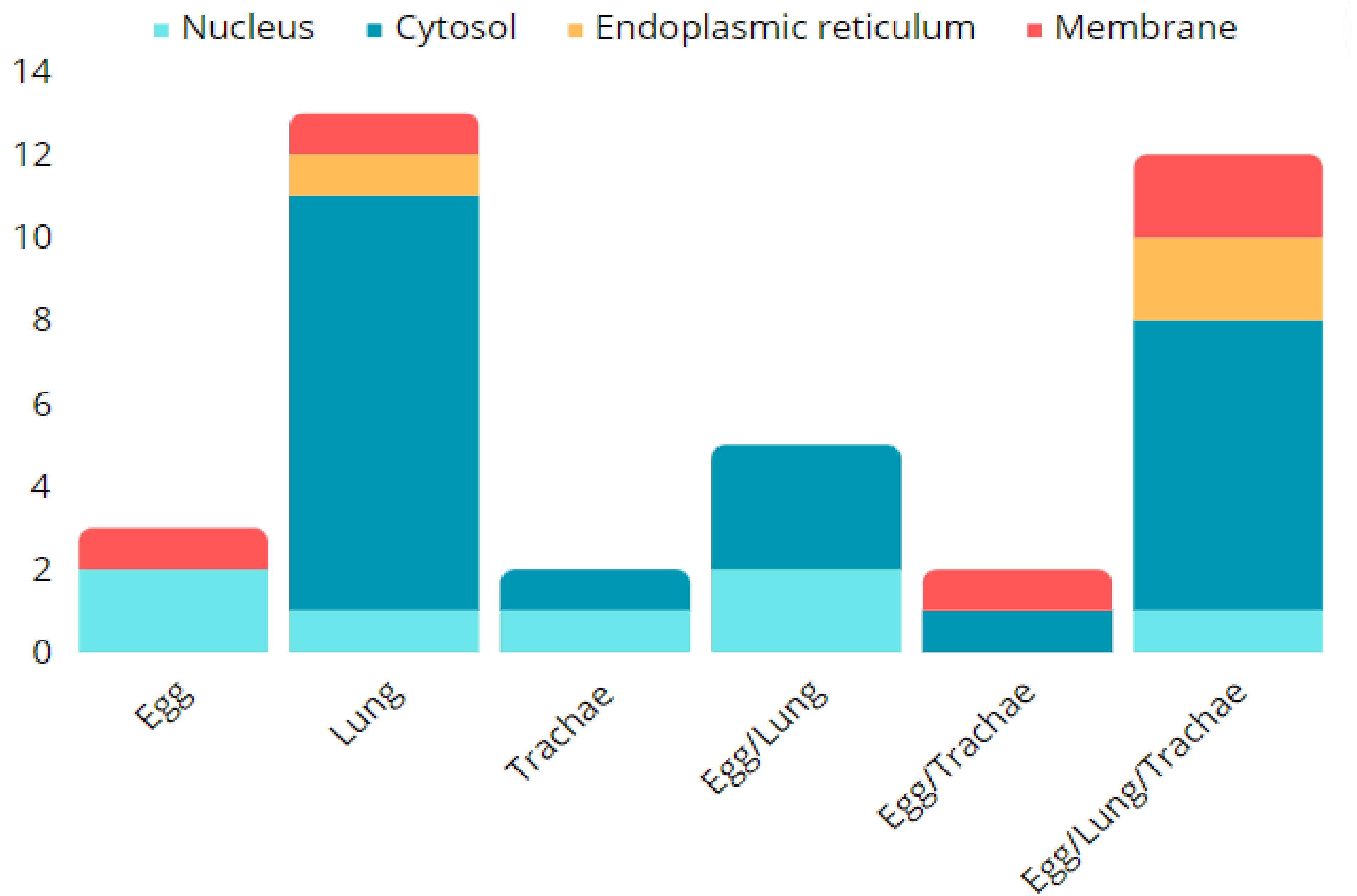

### Figure S2

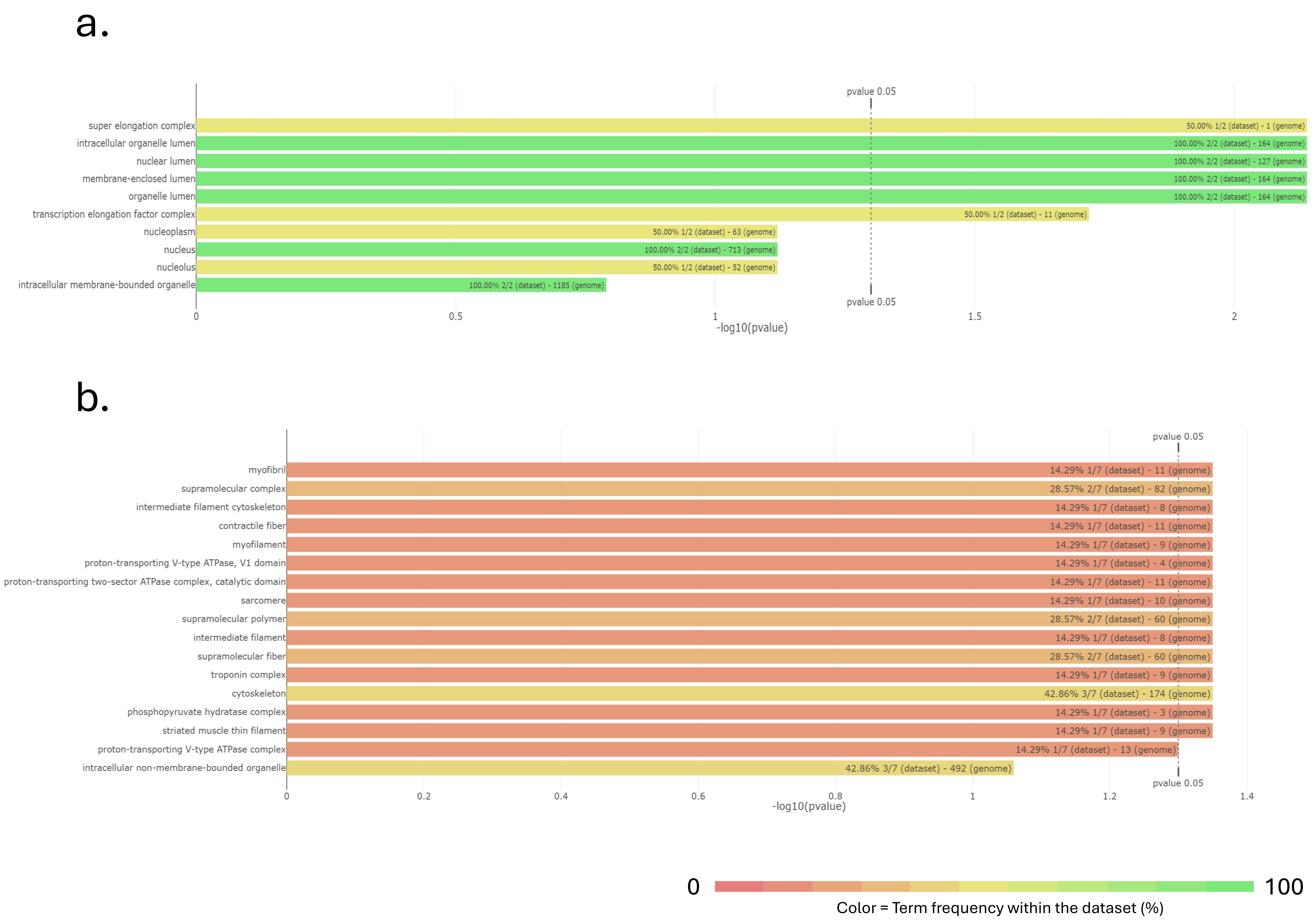

### Figure S3

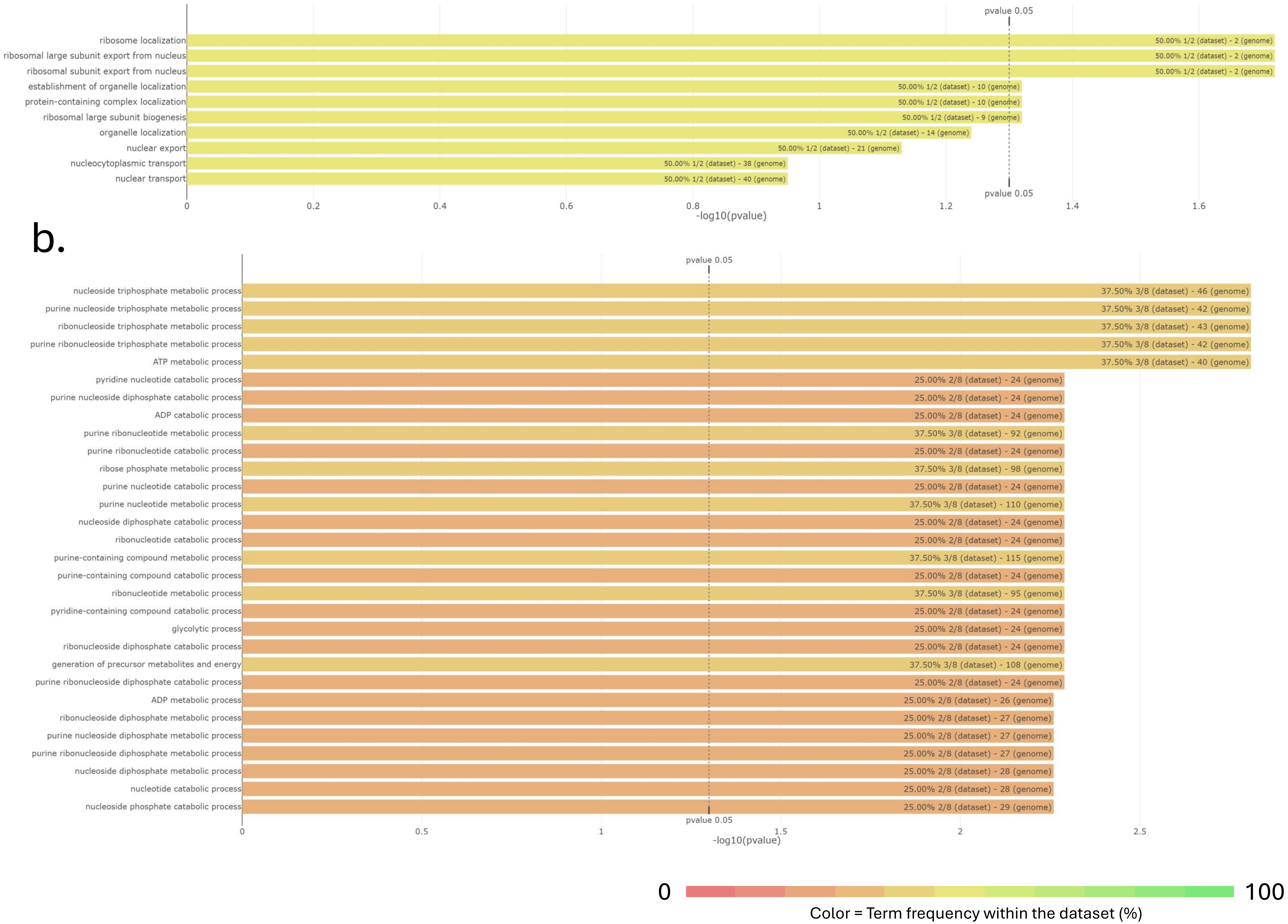
